## Supplemental Figures and Table Legends for "Extensive diversity in RNA termination and regulation revealed by transcriptome mapping for the Lyme pathogen *B. burgdorferi*"

[†]equal contribution

### Figure S1

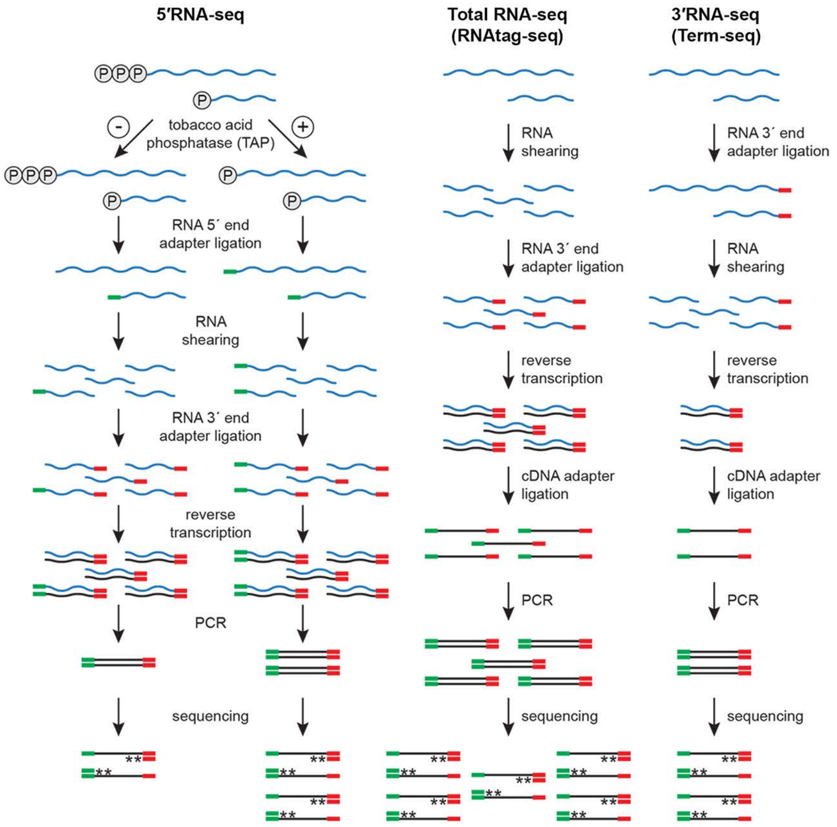
5′RNA-seq
Total RNA-seq
(RNAtag-seq)
3′RNA-seq
(Term-seq)
tobacco acid
phosphatase (TAP)
RNA 5´ end
adapter ligation
RNA
shearing
RNA 3´ end
adapter ligation
reverse
transcription
PCR
sequencing
RNA
shearing
RNA 3´ end
adapter ligation
reverse
transcription
cDNA adapter
ligation
PCR
sequencing
RNA 3´ end
adapter ligation
RNA
shearing
reverse
transcription
cDNA adapter
ligation
PCR
sequencing

**Figure S1**. RNA-seq approaches. Schematic of 5′RNA-seq (Adams et al., 2017b), total RNA-seq (modified from the RNAtag-seq methodology, (Shishkin et al., 2015)) and 3′RNA-seq (modified from the Term-seq methodology (Dar et al., 2016)). 5′ phosphates (circled ‘P’), RNA 5′ end adapter or cDNA adapter (green line), RNA 3′ end adapter (red line), and stranded sequencing (asterisks) are indicated.

### Figure S2

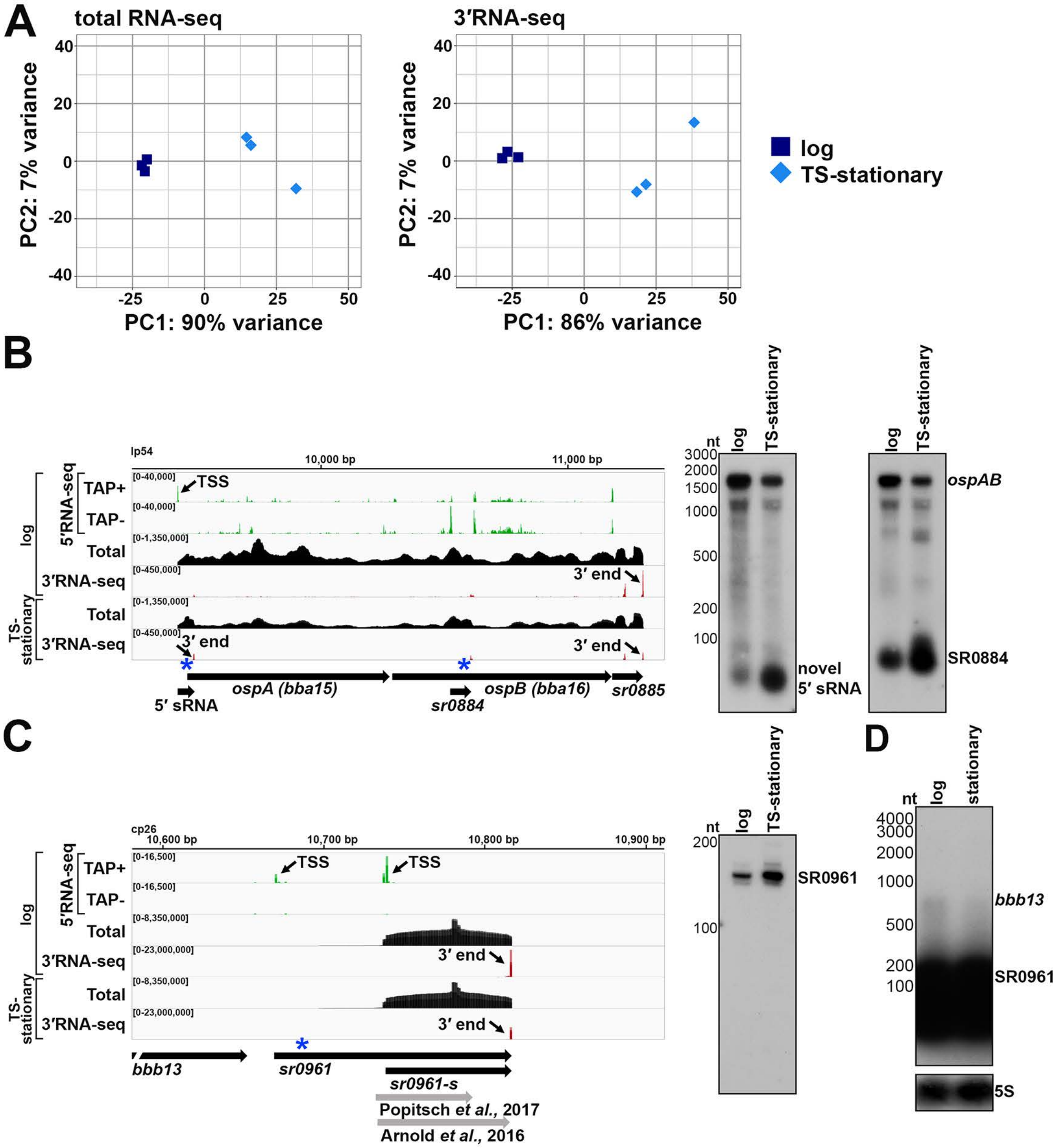
A
total RNA-seq
3′RNA-seq
PC2: 7% variance
PC1: 90% variance
PC1: 86% variance
log
TS-stationary
B
lp54
10,000 bp
11,000 bp
5′RNA-seq
TAP+
TAP-
TSS
Total
3′RNA-seq
3′ end
log
TS-stationary
5′ sRNA
ospA (bba15)
sr0884
ospB (bba16)
sr0885
nt
ospAB
novel 5′ sRNA
SR0884
C
cp26
10,600 bp
10,700 bp
10,800 bp
10,900 bp
bbb13
sr0961
sr0961-s
Popitsch et al., 2017
Arnold et al., 2016
SR0961
D
stationary
bbb13
5S

**Figure S2**. Analysis of 3′RNA-seq replicates and additional northern analyses of *ospAB* and SR0961. **(A)** Principal component analysis plot to show correlation among total RNA-seq and 3′RNA-seq replicates. For each sample, total RNA-seq read counts or 3’RNA-seq 3′ ends were quantified at annotated genes and normalized by variance-stabilizing transformation. Normalized counts were used in the principal component analysis to estimate the relationships between samples. RNA-seq browser image (left) and northern analysis (right) for the **(B)** *ospAB* locus and **(C-D)** SR0961 sRNA. Browser images display sequencing reads from logarithmic phase and TS-stationary phase cells, as in Figure 1. The northern analysis in panel B was performed on the same blot from Figure 1A; the northern analysis in panel C was performed on the same blot from Figure 1E; the northern analysis in panel D was performed on the same blot from Figure 5 and cropped to only show the logarithmic and stationary phase samples (RNAs were probed sequentially on the same membrane). The approximate probe sequence location is indicated by the blue asterisk on the corresponding browser image; **(D)** uses the same probe as in Figure 1E. Size markers are indicated for all RNAs. A previously unannotated fragment is denoted as ‘novel 5′ sRNA’.

### Figure S3

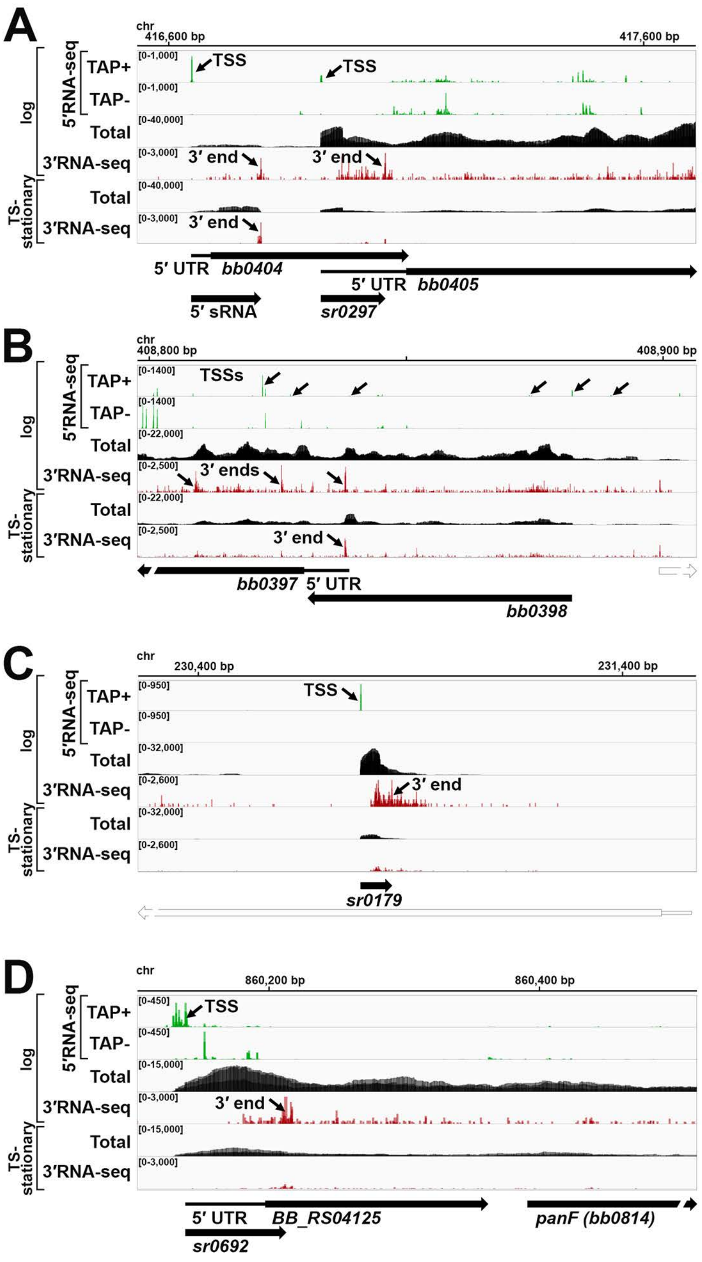

A
chr
416,600 bp
417,600 bp
log
5′RNA-seq
TAP+
TAP-
Total
3′RNA-seq
TS-stationary
Total
3′RNA-seq
TSS
TSS
3′ end
3′ end
3′ end
5′ UTR
bb0404
5′ UTR
bb0405
5′ sRNA
sr0297
B
408,800 bp
408,900 bp
TSSs
3′ ends
3′ end
bb0397
5′ UTR
bb0398
C
230,400 bp
231,400 bp
TSS
3′ end
sr0179
D
860,200 bp
860,400 bp
TSS
3′ end
5′ UTR
BB_RS04125
panF (bb0814)
sr0692

**Figure S3.** Examples of complex or ambiguous annotations. RNA-seq browser image for the **(A)** *bb0404-bb0405* locus, **(B)** *bb0398-bb0397* locus, **(C)** SR0179 locus, and **(D)** *BB_RS04125* and *panF* locus are shown. Browser images display sequencing reads from logarithmic phase and TS-stationary phase cells, as in Figure 1.

### Figure S4

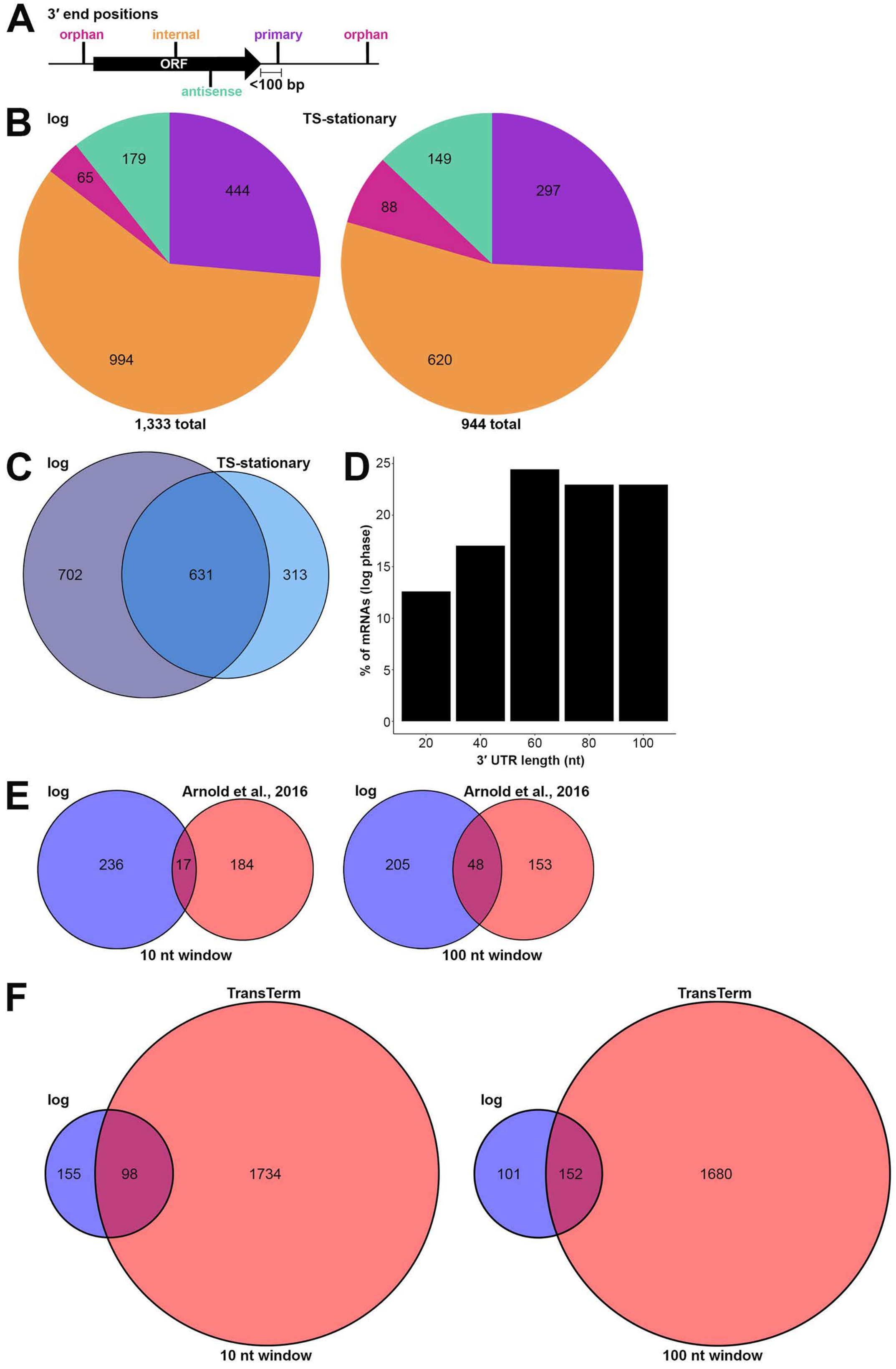
A
3′ end positions
orphan
internal
primary
orphan
ORF
antisense
<100 bp
B
log
179
65
444
994
1,333 total
TS-stationary
149
88
297
620
944 total
C
log
TS-stationary
702
631
313
D
% of mRNAs (log phase)
25
20
15
10
5
0
20
40
60
80
100
3′ UTR length (nt)
E
log
Arnold et al., 2016
236
17
184
10 nt window
log
Arnold et al., 2016
205
48
153
100 nt window
F
TransTerm
log
155
98
1734
10 nt window
TransTerm
log
101
152
1680
100 nt window

**Figure S4.** Distribution of 3′ ends and comparisons to other studies. **(A)** Schematic of classification of 3′RNA-seq identified ends relative to an annotated ORF. 3′ ends were defined as: primary (purple colored, located on the same strand within 100 bp downstream of the 3′ end of an annotated gene (mRNA ORF, tRNA, rRNA, or sRNA)), antisense (aquamarine colored, located on the opposite strand within 50 bp of a gene start and end coordinates), internal (orange colored, located on the same strand within a gene) and orphan (fuchsia colored, located in a 5′ UTR, long 3′ UTR or not falling in any of the previous classes). The black arrow represents an ORF. **(B)** Distribution of 3′RNA-seq detected ends relative to annotated genes for logarithmic and TS-stationary phase conditions. Some 3′ ends fit the criteria for two different categories; in the logarithmic phase condition: 100 are primary and antisense, 216 are primary and internal, 15 are internal and antisense, and 9 are primary, internal and antisense; in the TS-stationary phase condition: 64 are primary and antisense, 113 are primary and internal, 19 are internal and antisense, and 7 are primary, internal and antisense. **(C)** Comparison of 3′ ends identified in logarithmic phase culture to those identified in TS-stationary phase culture (Supplementary Table 1). If 3′ ends were called within a 10 nt window on the same DNA strand between both datasets, they were considered shared across those datasets. **(D)** Distribution of predicted 3′ UTR lengths for annotated mRNAs. The distance between the annotated stop codon and primary 3′ end for a total of 152 mRNAs are presented by intervals of 20 nt. **(E)** Comparison of current 3′RNA-seq to total RNA-seq data from Arnold et al. 3′RNA-seq, logarithmic phase condition. Detected 3′ ends with an intrinsic termination score ≥ 3.0 (Supplementary Table 1) were compared to previously predicted *B. burgdorferi* intrinsic terminators ((Arnold et al., 2016); Table S8, 'Predicted intrinsic terminators'). **(F)** Comparison of current 3′RNA-seq to terminators identified *in silico* by TransTerm. 3′RNA-seq, logarithmic phase condition. Detected 3′ ends with an intrinsic termination score ≥ 3.0 (Supplementary Table 1) were compared to predicted *B. burgdorferi* intrinsic terminators using the TransTerm terminator prediction algorithm (Kingsford et al., 2007). For panels E and F, if predicted terminators were called within a 10 or 100 nt window on the same DNA strand between both datasets, they were considered shared across those datasets.

#### Figure S5

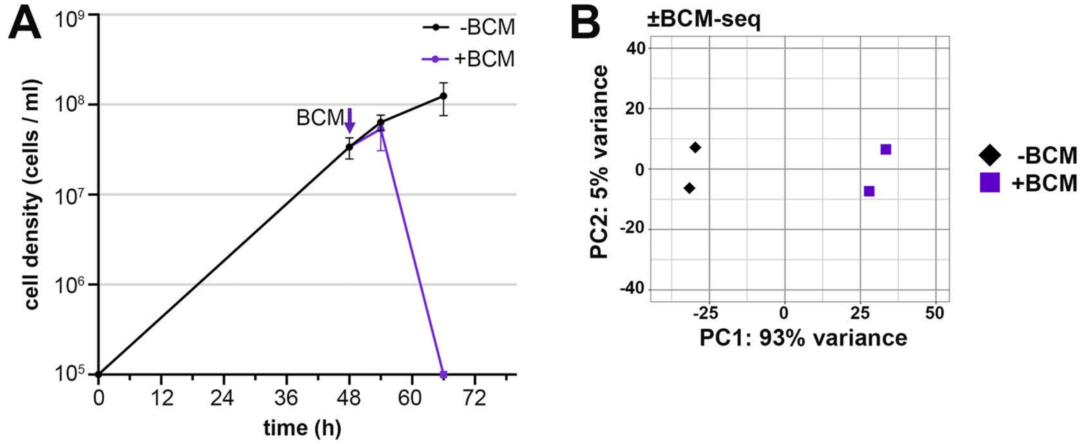
A
B
cell density (cells / ml)
time (h)
-BCM
+BCM
BCM
±BCM-seq
PC1: 93% variance
PC2: 5% variance
-BCM
+BCM

**Figure S5.** Effect of BCM treatment on *B. burgdorferi* growth and analysis of BCM-seq data. **(A)** Growth curve of ±BCM cultures. Cells (PA003) were grown to a density of $4.0x10^{7}$ cells/ml after dilution of the starter culture, the culture split and half treated with BCM for 6 h. Cell growth was monitored by dark field microscopy enumeration at the indicated time points. Each data point (circles) represents the average of two biological replicates (the same cultures used for the ±BCM-seq analysis), ± standard deviation. **(B)** Principal component analysis plot to show correlation among total -BCM and +BCM RNA-seq replicates. For each sample, read counts were quantified at annotated genes and normalized by variance-stabilizing transformation. Normalized counts were used in the principal component analysis to estimate the relationships between samples.

### Figure S6

**A** *bbk15*
*B. burgdorferi* B31 lp36: 9,118-9,999

TSS SR0821
1 ACATTTCAAAATATTGCAAAAAAATCATTTTTAAATTATATTTAATTTTATAATGATTACTATTTAATAT
NB Probe PA195
TTATTATAAGTTTGTATTATGGCTTTGCAGATATTCTGTAAAACCATTTTAAGGGCTTTATTTATGAAAA
AAAATTAATATTATTAATCAATATTTTTTTATTTATTTCATATCAAAACGATGGTTTAAATAAAAAATAAG
*bbk15* ORF
GCTAGCAAAAATCATAATTATAAAAAACAACTTGGGAAAAAATATGTGCGGGGAGAAAAGAATGAACTTT
281 TATCTTGGATAAATAATTTTGTAGAAGTTAATAATAAAGCCATAGATAATTCTTTTGAAAAATTCAAAAG
TGAGCCATCTAATCAATATAATCTTCCATTTAAAAATGTTAGTTGGTATACACTAAGGAAAACACTT...
841 ATAAATCTAGAAAAAGAGATTCGAAATATTATTGTTATTTAA

**B** *bgp (bb0588)*
*B. burgdorferi* B31 chromosome: 606,896 – 607,935

TSS SR0458
1 CAGCTGATAGCACTTGAACTAAATTCCCAAGATGAGGTATGTTATTAACATAGGGTAGAGCAGCTGTAAC
NB Probe PA145 TSS
TAGATTCATTTTTTTCATTCAGTTTTCCCTTTTTTTATGCTATTAATATTGTAATATGGATAGGATATTT
TTTTTCAATATCTTTTACAAAAATTCTTTCAAAAGCAATTCAATTTTGGCTTTTTTGTAAAAAATGATAT
*bgp* ORF
AAATATTCTTTTATTATAAGGAGTGTGATTTTATGAATAATTGTTTAATAAAGTTTTTTATTTTTTTATT
281 AGTTTTTTCAAACAGTTATGTTGCTTTTTCTAAAAATGTCAATGTTTTAATAGTAACTGCTATGGACTCT
GAGTTTGATCAGATAAATAAGCTTATGTCTAATAAGGAAGAAATAGTTCTTAAGGAGTATGGTCTTA...
981 TCTAAAATAGCTGCTTTCAATTCAGCCAAAGTTGTACAAGAAATTTTAAGAAAACTTTAA

**C** *bb0401*
*B. burgdorferi* B31 chromosome: 413,065-414,315

TSS SR0292
1 CTATTTATTATAAGTTTATATTTACAAAAAAATTTAGTACTATTTATTATACACAGTGTACCTTAAATCT
*bb0401* ORF
CTGAAAGGAGAAGTCATTATGAATATAAAAATCAATTTTTTTTTCACTTTGCCTATTGGAATCTTTTTAGG
NB Probe PA168
ATTGTTTTTCCCTCTTGGAATTTATAGCTCCTTATCACATGCTTTTATAAGATTATCATACTTATCTCTT
ATTCCCTTTTTAATATTTTCAATTCCATTAGGAATTGAAAATATTATTGAAAATAAAAACTTTAAAAAGC
281 TTTTTGGTAAAACAATTTATTATGGAATTTTAACTAACCTATCTGGAGTTGCTGTATCAATAATAGCTGC
AACAATATATCTTCCGCAAAGAATTCCAATACTAGAAAAAACAATACAAAATACATGTTTTTTTGAA...
1261 GAATTAAAAGATCAAGAAAAAATTAATTAA

**D** *oppAI (bb0328)*
*B. burgdorferi* B31 chromosome: 334,736-336,508

TSS SR0242
1 TTAAAATTATTAGACTTTTTTTATTATATAAGAATTAAAGTAAAATTTAAATCAAGACAAATCAAAGTAAGA
NB Probe PA083
CAAATCAAAGTACTGGAGGCATACATATTTAAAATTAAAACAAAATAAATATTCTTTATTAAAATTTATT
TSS *oppAI* ORF
AAAAAAGATTAATAAAGAATGAGAACAAAAAATATATAAAATATTGAAAAAGGAAAATCCCATGAAATAT
ATAAAAATAGCCTTAATGCTAATAATTTTTTCTTTAATAGCATGTATTAGTAATGCTAAAAAAGAAAAAA
281 TAGTTTTCAGAGTATCAAACTTAAGCGAGCCATCATCACTTGATCCTCAACTCTCAACAGACCTTTACGG
TAGCAACATTATTACAAACCTATTCTTAGGCCTAGCGGTAAAAGATTCTCAAACTGGAAAATATAAA...
1750 AAGATATTAAAACTAAAAAATAA

**E**

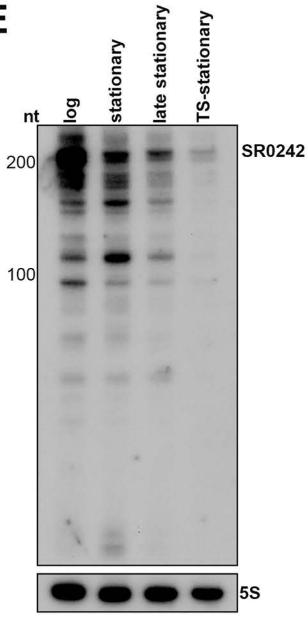


**F** *glpF (bb0240)*
*B. burgdorferi* B31 chromosome: 245,546 – 246,554

TSS
1 AATTCTTAAATTATTGACATTAATCTTAATTAAAAATAAGATATTAAATAATAATTTTAATAAGGCTTTT
NB Probe PA238
ATTAGAAAAATTAATTTTTTTTAATAAAAGAGACTAAAATAAAAAAATTCTAACCATCTTGCAAAAAACA
AAATAAAAATTTAATTAAAAGATTTAATATTATTCAGATTAAAAAATCAAAAATTAAAACTTCTTAATAAA
*glpF* ORF TSS
AAATAAAAATAGTTATAAAGATAAGGAGATATAATTATGAATTATACAAAATTCCAAGAATTTAATATCGGA
281 ATTTTTGGGAACATTTATCCTATTGGCTCTAGGAACTGGATCTGTTGCAATGACAGTATTATTTTCCTCA
AGTCCCGAAATACCAGGAGAAATAATAAAAGGAGGATATACAAATATAGTATTTGGATGGGGATTGGGTG
TAACGTTTGGTATTTACACAGCAGCAAGAATGAGCGGAGCACACCTAAACCCAGCTGTTAGCATAGG...
981 GAATTTACACTAAAAAAATAACAAAGACTAA

### Figure S6

**G** ***potAB (bb0640, bb0641)***
***B. burgdorferi*** **B31 chromosome: 681,839 – 679,754**

$P_{potA}$
1 CACTTGATATTCTACAAAATTTTGCAAAAGCAAGAAAATTAATCGGTAAAAAAAATGAACTTAACCTTGA
AAAAGCATCAAAAATATTAATCAAAGAATTTAGAGAGGGTAAATTTGGCAAAATAATTCTTGATAAGAAT
TSS
TATAATGCCTTTTAAAAAAGGCATTTACATAAATAATAATATTAAGTATAATCTTGATTTGTATTAATAC
*potA* ORF NB Probe PA102
AGCCATAAGGAGGTTGGAATTAGTTGGATAATTGTATCCTAGAGATTAAAAATCTAAGTCATTATTATGA
281 TAACAATGGAAACAAAACTTTAGATAACATAAATTTAAAAATTAAAAAAAATGAGTTTATCACACTACTA
GGCCCATCCGGATGTGGAAAAACAACATTGATAAAAATATTGGGTGGTTTTTTAAGCCAAAAAAATGGAG
AAATTTATTTCTTTTCTAAAGAAATATCTAAAACCAGTCCAAACAAAAGAGAAATTAATACTGTATTTCA
AAATTATGCACTTTTCCCACATATGAATGTTTTTGACAATATTTCATTTGGACTTAGAATGAAAAAAACG
561 CCAAAAGATATAATCAAAGAAAAAGTAAAAACATCGCTTTCGCTGATAGGAATGCCAAAATACGCATACA
$P_{potB}$
GAAATATTAACGAACTATCGGGGGGGCAAAAGCAAAGAGTTGCAATAGCAAGAGCAATGGTAATGGAACC
TAAGCTTTTACTCCTAGATGAACCACTTTCCGCGCTTGATTTGAAAATGCGACAAGAGATGCAAAAAGAA
TTAAAAAAAATACAGCGTCAGCTTGGAATCACATTCATATATGTTACTCACGATCAAGAAGAGGCATTGA
TSS *potB* 5'$_{1-561}$
841 CAATGAGTGACAGAATCGTTGTAATGAATGAAGGAATAATTCTGCAAATAGGAACACCTGAGGAAATTTA
CAATGAGCCTAAAACAAAGTTTGTAGCCGATTTTATTGGAGAAAGCAATATTTTTGATGGAACATATAAA
AAAGAGCTGGTTGTAAGTTTGCTTGGTCATGAATTTGAATGCCTTGACAAAGGATTTGAAGCTGAAGAAG
CAGTTGACCTTGTAATACGCCCAGAAGATGTAAAACTACTTCCAAAAGGAAAAGGACATTTAAGCGGAAC
1121 TATAACATCAGCAATTTTTCAAGGAGTTCATTACGAAATGACTCTAGAAATCCAAAAAACAAATTGGATA
*potB* 5'$_{334-561}$
GTTCAAAGCACAAGACTTACAAAAGTTGGAGAAGAAGTTGATATATTTTTAGAACCTGATGATATTCATG
*potB* ORF
TTATGCATAAGGAATAATGGTTTTGAAAAAGTTGATATTAATCATATACTCCATATTCCTACTAACATTT
*potB* 5'$_{485-561}$ (SR0541) NB Probe PA086
AGTATTCTTCCCTTACTAATAATAATATTGCTTGGATTTTTAAATGAAAAAAACGAATTTACCATCTATA
1401 ATTTCATTGGACTTTTAAATCCAAGCTATCTTAATATTTTTTCAAGAAGTCTAAAACTCGCAACAATAGC
NB Probe PA093
AACAATTTTTTGCATTTTAATAGGCTATCCTGCCGCTTGGCTAATTTCATTATCAAAAAAAAGTGCTCAA
AACAAATTAATAATCATGATAATACTTCCTATGTGGATAAATACATTACTTAGAACTTATGCCTGGATGA
GAATACTTGGAAAAAAACGGATTCATCAACAACTTATTTGAAAAGATCGGAATTGGAACTTTAGATCTTCT
1680 TTATAATGAACAGGCTGTTACAATAGGCATGATATACAATTTTTTGCCTTTTATGATCTTGCCAATATAC
ACGGGGCTTTTAAAAATTAAGCCAGAATATATTGAAGCATCACAAGATCTTGGAGCAAGAATGTGGCAAA
TATTACTTTATATAAAAATACCACTAACACTCTCTTACCTGGCAACAGGAATAATTATGGTATTTATTCC
TTCAATTACGGTATTTATCATTTCAGATTTGCTAGGAGGCTCTAAACAAATTTTAATAGGAAATCTAATA
1959 AGCAAACAGTTTCTCTTTATAGAAGACTGGAATACTGGGGCTGCAATTTCATTTATTTTAATGTTAGTAA
TATTAATTTTTAATTTAATAATAATAAAATTAATGCGAAAAAATAATGGGGAGTAA

**H**

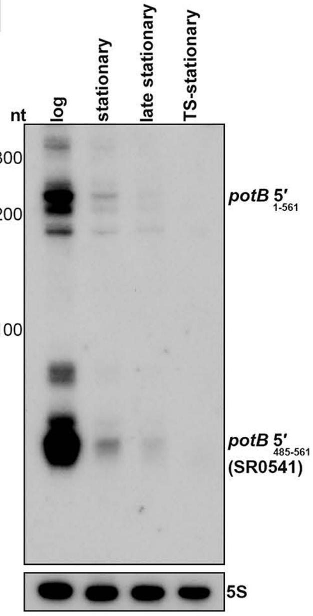


**I**

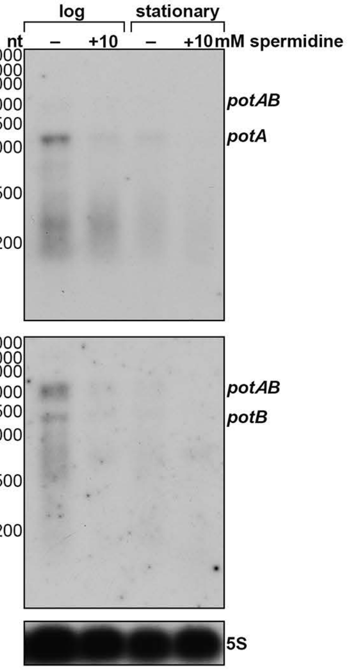

**Figure S6.** 5′ mRNA sequences that harbor 3′ ends. DNA sequences for the **(A)** *bbk15* locus, **(B)** *bgp* locus, **(C)** *bb0401* locus, **(D)** *oppAI* locus, **(F)** *glpF* locus and **(G)** *potAB* locus are shown. Chromosome coordinates are indicated above the sequence for each locus. TSS(s) identified by 5′RNA-seq (Adams et al., 2017b) are highlighted in green and 3′ ends identified by 3′RNA-seq, logarithmic phase condition (Supplementary Table 1), are highlighted in red. Predicted regions of 5′ mRNA fragments, based on RNA-seq identified 5′ and 3′ ends and band sizes from the northern analysis (Figure 5), are highlighted in grey. Annotated start and stop codons are labeled with green and red text, respectively. Probe sequences used for the northern analysis in Figure 5 are highlighted in yellow. For panel G, *potA/B* promoter sequences used for luciferase transcriptional fusions in Figure 7 are underlined with a wavy line, and the *potB* sequence used for luciferase translational fusions (*potB*$_{1\text{-}594}$) is underlined with a solid line. Northern analysis of SR0242 (**E**) and potB 5′ (**H**). Northern analysis was performed using the same RNA analyzed by agarose northern analysis in Figure 5. Total RNA was separated on an acrylamide gel, transferred to a membrane and probed for the RNAs, sequentially on the same membrane. The 5S blot in panel E was repeated in panel H. **(I)** Northern analysis of effects of spermidine on *potAB*; performed on the same blot from Figure 6E (RNAs probed sequentially on the same membrane). The 5S blot in panel I was repeated from Figure 6E.

### Figure S7

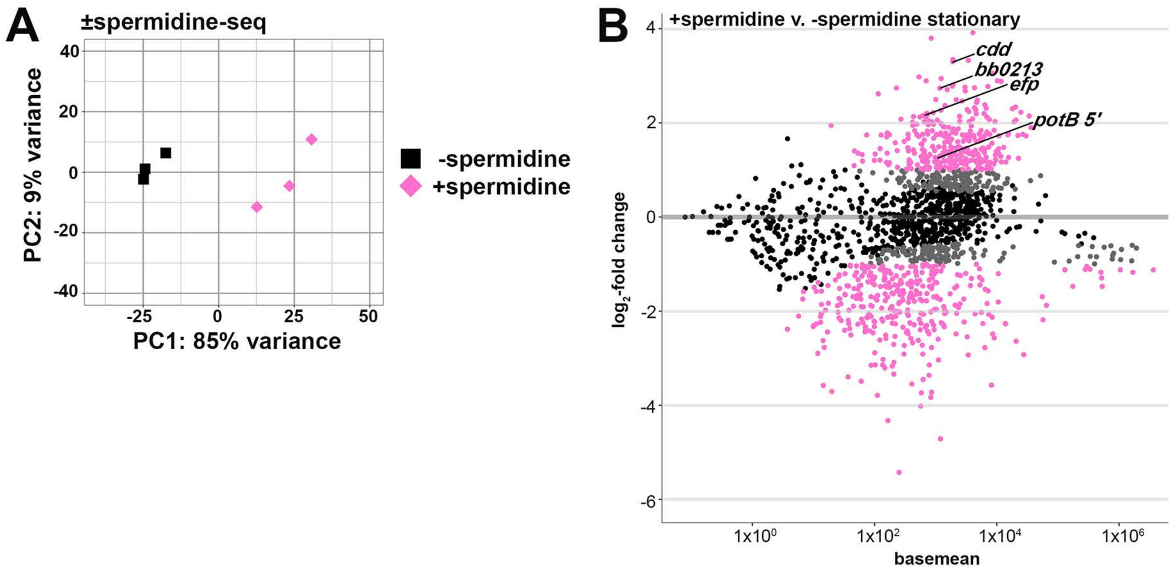
A
±spermidine-seq
PC2: 9% variance
PC1: 85% variance
40
20
0
-20
-40
-25
0
25
50
-spermidine
+spermidine
B
+spermidine v. -spermidine stationary
cdd
bb0213
efp
potB 5'
log2-fold change
basemean
4
2
0
-2
-4
-6
1x10^0
1x10^2
1x10^4
1x10^6

**Figure S7**. Spermidine globally affects RNA levels. **(A)** Principal component analysis plot to show correlation among -spermidine and +spermidine total RNA-seq replicates. For each sample, read counts were quantified at annotated genes and normalized by variance-stabilizing transformation. Normalized counts were used in the principal component analysis to estimate the relationships between samples. **(B)** MA-plot outputting results from DESeq2 analysis, comparing RNA levels in cells grown with/without the addition of 10mM spermidine in BSK-II. Three independent biological repeats of WT (PA003) were grown to stationary phase (as described for Figure 6E) prior to cell lysis, RNA isolation and sequencing. Data are plotted by average $\log_2$-fold change (LFC) of annotated ORFs and sRNAs for +spermidine versus -spermidine (y-axis) over average normalized read count values ('basemean', x-axis). Black points are non-significant ($p>0.01$), grey values are significant but have a LFC $< 1.0$, and pink points are significant with a LFC $\geq 1.0$.

### Figure S8

**A**

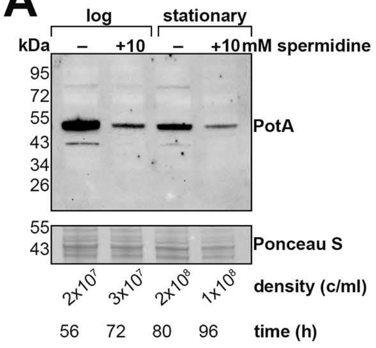


**B**

***potB*** $_{1-561}$
***B. burgdorferi* B31 chromosome: 680,968 – 680,388**

TSS ***potB* 5'** $_{1-561}$

```
1   AGGAATAATTCTGCAAATAGGAACACCTGAGGAAATTTACAATGAGCCTAAAACAAAGTTTGTAGCCGATTT
    TATTGGAGAAAGCAATATTTTTGATGGAACATATAAAAAAGAGCTGGTTGTAAGTTTGCTTGGTCATGAATT
    TGAATGCCTTGACAAAGGATTTGAAGCTGAAGAAGCAGTTGACCTTGTAATACGCCCAGAAGATGTAAAACT
217 ACTTCCAAAAGGAAAAGGACATTTAAGCGGAACTATAACATCAGCAATTTTTCAAGGAGTTCATTACGAAAT
    GACTCTAGAAATCCAAAAAACAAATTGGATAGTTCAAAGCACAAGACTTACAAAAGTTGGAGAAGAAGTTGA
                                                      potB ORF
    TATATTTTTAGAACCTGATGATATTCATGTTATGCATAAGGAATAATGGTTTTGAAAAAGTTGATATTAATC
                                                      LeuLysLysLeuTyrLeuIle
                                                       potB 5'485-561 (SR0541)
433 ATATACTCCATATTCCTACTAACATTTAGTATTCTTCCCTTACTAATAATAATATTGCTTGGATTTTTAAAT
    IleTyrSerIlePheLeuLeuThrPheSerIleLeuProLeuLeuIleIleIleLeuLeuGlyPheLeuAsn
    GAAAAAAACGAATTTACCATCTATAATTTCATTGGACTTTTAAATCCAAGCTATCTTAATATTTTTCAAGA
    GluLysAsnGluPheThrIleTyrAsnPheIleGlyLeuLeuAsnProSerTyrLeuAsnIlePheSerArg
    AGTCTAAAACTCGCAACA 594
    SerLeuLysLeuAlaThr
```

**C**

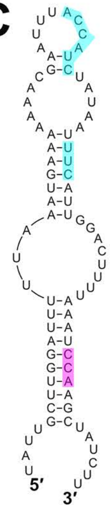


**SR0541**

**Figure S8.** PotA levels decrease with spermidine exposure and features of the *potB*$_{1\text{-}561}$ sequence. **(A)** Western analysis of PotA levels. Cells were grown in the presence of 10 mM spermidine as in Figure 6D. Samples were harvested at the indicated time points to best match cell density (indicated below each sample) and total protein isolated. Protein extracts were separated on a Tris-Glycine gel, stained using Ponceau S stain, and probed with α-PotA antibodies (Lin et al., 2017). **(B)** DNA sequence of *potB*$_{1\text{-}594}$ (used for translational luciferase fusions), as in Figure S6G. Amino acid sequence of PotB is included beneath the corresponding DNA sequence, with rare codons highlighted in blue and proline residues highlighted in magenta. The PotA stop codon is indicated with red text. **(C)** Structure of the *potB*$_{1\text{-}561}$ RNA as modeled by RNAfold Web Server (Gruber et al., 2008). Nucleotide sequences encoding rare codons and proline residues are highlighted in blue and magenta respectively, as in panel B.

#### SUPPLEMENTAL TABLE LEGENDS

**Supplementary Table 1.** 3′ ends identified by 3′RNA-seq. Curated 3′ ends for *B. burgdorferi* B31 grown to logarithmic phase at 35°C or temperature-shifted (~23°C to 35°C) stationary phase. The data for each growth condition are displayed on a separate tab. The replicon, genomic coordinate of the 3′ end ('3′ end position'), DNA strand, average RNA-seq read count of the 3′ end of biological triplicates, 3′ end classification (see Materials and Methods), the gene ID, NCBI locus tag, and gene name(s) for the classification ('details'), the gene descriptions, the sequence surrounding the 3′ end (50 bp upstream and 10 bp downstream, 3′ end nucleotide red and bolded), and the distance of a primary 3′ end from the upstream mRNA ('predicted 3′ UTR length') make up the columns of the table. Each 3′ end was also given an intrinsic terminator score (Chen et al., 2013), which includes an output of the Kinefold structure (Xayaphoummine et al., 2005) with possible A-tract, hairpin, loop and U-tract. Intrinsic terminator scores ≥ 3.0 are suggestive of intrinsic termination. An 'undefined' intrinsic terminator score indicates one that could not be calculated because the sequence could not be folded into a recognizable secondary structure. An 'impossible' intrinsic terminator score indicates one that is too small to be accurately calculated.

**Supplementary Table 2.** Previously annotated *B. burgdorferi* sRNAs with defined 5′ and 3′ ends. The sRNA ID, based on the nomenclature from (Popitsch et al., 2017), replicon, genomic coordinates as determined algorithmically in this study ('annotated 5′ coordinate' and 'annotated 3′ coordinate'), and DNA strand are listed for each sRNA. Additional columns give the 5′ most coordinate, 5′ coordinate with the highest read count, possible TSS(s) and possible processed 5′ end(s) determined by visual inspection of 5′RNA-seq (Adams et al., 2017b) as well as possible 3′ end(s) determined by visual inspection of 3′RNA-seq (this study). The dataset

((Popitsch et al., 2017) or (Arnold et al., 2016)) where the sRNA was previously detected is indicated. See Materials and Methods for details of visual inspection.

**Supplementary Table 3.** Updated *B. burgdorferi* B31 gene annotations. The replicon and its NCBI reference number, DNA strand, genomic coordinate of the 5′ and 3′ ends, sequence length, classification, commonly used gene IDs (‘gene ID’), NCBI locus tag, and gene name(s) with gene descriptions from NCBI and UniProt make up the columns of the table. 5′ and 3′ UTR regions are predicted algorithmically using mRNA primary ends identified from 5′RNA-seq (Adams et al., 2017b) and 3′RNA-seq (this study), respectively. Characterized and/or identified genes not listed in NCBI or UniProt databases are also noted.

**Supplementary Table 4.** Identification of Rho termination regions using ±BCM-seq. The data for each replicate are displayed on a separate tab. All identified Rho termination regions are represented, defined as regions with at least one genomic coordinate with a significance score $<1e^{-4}$. Rho scores were calculated for each genomic position by comparing ±BCM-seq coverage in windows 800 nt upstream and downstream in the treated (+BCM) and untreated (-BCM) samples (see Materials and Methods). The replicon, genomic coordinate with the significant highest Rho score within the 800 nt window (‘Rho region position’), DNA strand, Rho region position classification (using the same parameters for classifying 3′ ends; see Materials and Methods), the gene ID and name(s) for the classification (‘details’), the read coverage in the 800 nt windows upstream and downstream the Rho region position ±BCM, the Rho score, and the *p*-value from the Fisher’s exact test (‘significance score’) make up the columns of the table. An ‘undefined’ Rho score indicates one that could not be calculated due to an absence of reads in the ±BCM adjacent regions. A significance score of ‘n/a’ indicates that the significance score was too low ($< 1e^{-300}$) to accurately report. The third tab in the table summarizes, for each gene, if a Rho region was found 800 nt downstream of or internal to all annotated ORFs/sRNAs for either, both or neither replicate dataset.

**Supplementary Table 5.** Analysis of 3′ ends in 5′ UTRs and within coding sequences. 3′RNA-seq identified 3′ ends for *B. burgdorferi* B31 that were between 200 nt upstream of an ORF and the corresponding stop codon. The replicon, genomic coordinate of the 3′ end ('3′ end position'), DNA strand, average RNA-seq read count of the 3′ end of biological triplicates, 3′ end classification (see Materials and Methods), the gene ID for the classification ('details'), the location of the 3′ end relative to an ORF – upstream ORF or internal ('ORF classification'), the distance of the 3′ end from the gene's annotated start codon, and the gene annotation of the associated ORF ('upstream ORF/internal details') make up the columns of the table. Each 3′ end was also given an intrinsic terminator score (see Materials and Methods), Rho score (an assessment of whether it is in a Rho termination region), and spermidine-dependent score (an assessment of whether its generation is impacted by spermidine treatment, see Materials and Methods). Intrinsic terminator scores ≥ 3.0 are suggestive of intrinsic termination. An 'undefined' intrinsic terminator score indicates one that could not be calculated because the sequence could not be folded into a recognizable secondary structure. Rho and spermidine-dependent scores ≥ 2.0 are suggestive of Rho termination and spermidine-affected, respectively. An 'undefined' Rho score indicates one that could not be calculated due to an absence of reads in the ±BCM adjacent regions. An 'undefined' spermidine-dependent score indicates one that could not be calculated due to an absence of reads in the ±spermidine adjacent regions. A significance score of 'n/a' indicates that the Rho or spermidine-dependent significance score was too low (< $1e^{-300}$) to accurately report. No significance scores were calculated for 'undefined' scores.

**Supplementary Table 6.** RNA-seq analysis of transcript levels after exposure to spermidine. Differential expression analysis was carried out using DESeq2 with the annotated ORFs and sRNAs listed in Supplementary Table 3. The replicon, DNA strand, the gene ID, NCBI/UniProt locus tag, and gene name(s) for the classification ('details') and gene descriptions, basemean, $log_2$-fold change, lfcSE (standard error of the $log_2$-fold change) and adjusted p-value ('padj') make up the columns of the table. DESeq2 analysis was performed in two ways, one which included reads overlapping 2 or more features (columns labelled with the suffix (allowOverlap)), one performed with standard quantification parameters that included only reads overlapping a single feature. A basemean value of 0 indicates the absence of reads in the feature in either condition. No $log_2$-fold change, lfcSE, or adjusted p-value were calculated for features with a basemean of 0.

**Supplementary Table 7.** Raw data for luciferase assays (tab 1) and RT-qPCR (tab2). Bolded values in the table represent the plotted datapoints in Figure 7.

**Supplementary Table 8.** List of strains together and plasmids (tab 1) as well as oligonucleotides (tab 2) used in this study.
